# The interactome of site-specifically acetylated linker histone H1

**DOI:** 10.1101/2021.06.08.447242

**Authors:** Eva Höllmüller, Katharina Greiner, Simon M. Kienle, Martin Scheffner, Andreas Marx, Florian Stengel

## Abstract

Linker histone H1 plays a key role in chromatin organization and maintenance, yet our knowledge of the regulation of H1 functions by posttranslational modifications (PTMs) is rather limited. In this study, we report on the generation of site-specifically mono- and di-acetylated linker histone H1.2 by genetic code expansion. We used these modified histones to identify and characterize the acetylation-dependent cellular interactome of H1.2 by affinity purification-mass spectrometry (AP-MS) and show that site-specific acetylation results in overlapping, but distinct groups of interacting partners. Among these, we find multiple translational initiation factors and transcriptional regulators such as the NAD^+^-dependent deacetylase SIRT1, which we demonstrate to act on acetylated H1.2. Taken together our data suggests that site-specific acetylation of H1.2 plays a role in modulating protein-protein interactions.

## INTRODUCTION

Linker histone H1 proteins are small, highly basic proteins and represent one of the five histone protein families. Histones are responsible for the condensation and organization of DNA within the nucleus of all eukaryotes. The DNA is wrapped around an octamer of core histones (H2A, H2B, H3, H4) to form nucleosomes and H1 proteins additionally bind at the DNA entry and exit sites to form the chromatosome. Thus, linker histones mediate the formation of higher-order chromatin structures and both stabilize nucleosomes and provide chromatin with structural and functional flexibility. Human cells encode eleven H1 variants including seven somatic subtypes (H1.1-H1.5, H1.0, H1.x) with H1.2 and H1.4 being the two predominant variants. H1 proteins share a tripartite structure, consisting of a central globular domain (GD) flanked by long, unstructured N-terminal (N-terminal domain, NTD) and C-terminal (C-terminal domain, CTD) tails.^1–6 (and references therein)^

Histones are subjected to a diverse set of posttranslational modifications (PTMs) including acetylation, methylation, and ubiquitylation. These modifications generate a complex epigenetic code, which modulates protein-protein and protein-DNA interactions, resulting in the regulation of chromatin structure and function, a concept termed the ‘histone code’. To decipher the functional relevance of certain histone modifications, their specific effects need to be determined.^7–12^ Although the H1 modification pattern appears to be as complex as that of core histones, relatively little is known about the effect of PTMs on H1 function. Obstacles in the elucidation of the role of H1 PTMs include H1 subtype heterogeneity, different modifications between different H1 subtypes, and the lack of site- and modification-specific antibodies.^2, 3, 5, 6^ Furthermore, intrinsic characteristics of H1, as the long, highly unstructured and lysine-rich CTD, which is prone to degradation and yields insoluble, truncated proteins,^13, 14^ have hampered even *in vitro* studies until recently.^15^ In addition to its role as a general organizer of chromatin architecture,^1^ there is growing evidence that H1 regulates cellular functions also *via* direct protein-protein interactions.^15–19^ Along this line, acetylation of linker histones has been linked to activation of gene expression^20^ and shaping of chromatin architecture and dynamics.^21, 22^ For example, it was shown that H1.4 acetylated at position K34 is enriched at promoters of active genes and results in the recruitment of a general transcription factor (TAF1 subunit of transcription factor TFIID),^20^ while a study using an H1 variant using an acetylation-mimetic amino acid at position K85 (H1 K85Q) observed chromatin condensation and facilitated chromatin compaction.^22^

To unravel the influence of specific PTMs on the histone code, the generation of defined, site-specifically modified histones by chemical protein synthesis and chemical biology methods has been proven extremely powerful.^11^ Methods include cysteine bioconjugation,^23, 24^ protein semisynthesis,^25–28^ and genetic code expansion.^29, 30^ Recently, we showed that the generation of ubiquitylated H1.2-conjugates using selective pressure incorporation and amber codon suppression in combination with Cu(I)-catalyzed azide-alkyne cycloaddition (CuAAC, click reaction) followed by proteomic profiling represents an attractive approach to study potential effects of PTMs on linker histone H1 function.^15^ Here, we have expanded this approach to generate site-specifically, homogeneously mono- and diacetylated H1.2 by amber codon suppression. In combination with affinity purification-mass spectrometry (AP-MS), this allowed us to identify acetylation-specific interactors of H1 and to demonstrate the role of this PTM in modulating the cellular H1 interactome.

## MATERIAL AND METHODS

### Expression and purification of H1.2 and H1.2 KxAc

H1.2 was expressed and purified as described.^15^ In brief, the cDNA encoding human H1.2 with an N-terminal His_6_-tag and a C-terminal Strep-tag II was inserted into the multiple cloning site of pET11a which additionally contained the expression cassette for the tRNA_CUA_ gene in its backbone. For incorporation of acetyllysine (AcK) by amber stop codon suppression, the K17, K64, or both (K17K64) codons of H1.2 were replaced by an amber stop codon (TAG/UAG) using site-directed mutagenesis. *Escherichia coli* (*E. coli)* BL21 (DE3) cells were transformed with these plasmids and the pRSFDuet-1 vector, containing the pyrrolysine tRNA synthetase gene with mutations in the active site of the enzyme as previously described.^31^ Cells were cultured in LB medium supplemented with carbenicillin and kanamycin at 37 °C. At OD_600_ = 0.3, 10 mM (K17, K64) or 15 mM (K17K64) AcK (abcr) and 20 mM nicotinamide (Sigma-Aldrich) were added. Gene expression was induced at OD_600_ = 0.7 by the addition of 1 mM IPTG. After 6 h, cells were harvested by centrifugation. For expression of H1.2, only the respective pET11a plasmid was transformed into *E. coli* BL21 (DE3) and neither AcK nor nicotinamide were added during expression.

To isolate H1.2 or H1.2 KxAc from inclusion bodies, cells were lysed by sonication in 50 mM Tris-HCl (pH 8.5), 100 mM NaCl, 1 mM EDTA, 1% (v/v) Triton X-100, 2 mM PMSF. Pellets were incubated in 50 mM Tris-HCl (pH 7.0), 1 M NaCl, 6 M urea, 10 mM β-mercaptoethanol. After centrifugation, cOmplete His-Tag Purification Resin (Roche) was added to the supernatant and samples were incubated overnight at 4 °C. Beads were washed with increasing concentrations of imidazole and finally eluted with up to 500 mM imidazole. Elution fractions were pooled, dialyzed in water, and concentrated by ultrafiltration. Protein concentration was determined by BCA assay (Thermo Fisher Scientific) and SDS-PAGE analysis followed by Coomassie blue staining.

Incorporation of AcK was verified by LC-MS/MS analysis. Proteins were separated by SDS-PAGE, visualized with Coomassie blue staining, and the respective bands prepared for analysis by in-gel digestion.^32^ In short, gel pieces were destained in 50 mM NH_4_HCO_3_, 50% (v/v) acetonitrile (ACN) and washed with 50 mM NH_4_HCO_3_. Disulfide bonds were reduced by the addition of 10 mM DTT in 50 mM NH_4_HCO_3_ for 60 min at 56 °C, followed by alkylation in 50 mM 2-iodoacetamide in 50 mM NH_4_HCO_3_ for 60 min at room temperature in the dark. After washing and dehydration in ACN, proteins were digested overnight at 37 °C using trypsin (1:50, w/w, Promega). Peptides were extracted in ACN and formic acid and combined extracts were freeze-dried. Before LC-MS/MS analysis, samples were desalted using Micro-ZipTips (μ-C18, Merck).

Peptides were analyzed on a QExcactive HF Hybrid Quadrupole-Orbitrap (Thermo Fisher Scientific) coupled to an EASY-nLC 1200 system. The gradient (flow rate of 300 nl/min) for peptide separation started from 2.5% ACN, 0.1% formic acid to 32% ACN, 0.1% formic acid in 40 min, followed by a washing step at 75% ACN. The mass spectrometer was operated in data-dependent Top20 mode with dynamic exclusion set to 40 s. Full scan MS spectra were acquired at a resolution of 120,000 (at m/z 200), scan range 350-1600 m/z with an automatic gain control target value of 3e^6^ and a maximum injection time of 60 ms. Most intense precursors with charge states of 2-6 reaching a minimum automatic gain control target value of 1e^3^ were selected for MS/MS experiments. Normalized collision energy was set to 28%. MS/MS spectra were collected at a resolution of 30,000 (at m/z 200), an automatic gain control target value of 1e^5^ and 50 ms maximum injection time.

Raw files were analyzed using the Mascot 2.5.1 search engine MS/MS ions search (Matrix Science). Oxidation (M) and Acetylation (K) were set as variable modifications and trypsin cleavage allowing up to two missed cleavages was chosen.

For determination of the mass of full-length proteins, recombinant proteins were analyzed on a micrOTOF II (Bruker) coupled to an Agilent 1200 HPLC system. Compass DataAnalysis (Bruker) software was used for re-calibration and spectra deconvolution.

### Nucleosome and chromatosome assembly

Nucleosomes were assembled by stepwise salt dialysis. DNA was prepared by PCR using the *Widom 601* DNA sequence as template^33^ 5’-CTATACGCGGCCGCCCTGGAGAATCCCGGTGCCGAGGCCGCTCAATTGGTCGTAGCAAGCTCTAGCACCGCTTAAACGCACGTACGCGCTGTCCCCCGCGTTTTAACCGCCAAGGGGATTACTCCCTAGTCTCCAGGCACGTGTCAGATATATACATCCTGTGCATGTGGATCCGAAT-3’ with 5’-CTATACGCGGCCGCCCTGG-3’ and 5’-desthiobiotin-TEG-ATTCGGATCCACATGCACAGGATG-3’ as primers. Finally, the DNA was purified by EtOH precipitation. 100 ng/μl *601* DNA, 40 ng/μl recombinant chicken histone octamer (Abcam), 6 ng/μl pUC18 as competitor DNA, and 200 ng/μl BSA were mixed and dialyzed at 4 °C in high salt buffer 10 mM Tris-HCl (pH 7.6), 2 M NaCl, 1 mM EDTA, 0.05% (v/v) Nonidet P-40 Substitute, 1 mM β-mercaptoethanol, followed by stepwise reduction of the NaCl concentration to end up in low salt buffer 10 mM Tris-HCl (pH 7.6), 50 mM NaCl, 1 mM EDTA, 0.05% (v/v) Nonidet P-40 Substitute, 1 mM β-mercaptoethanol. For chromatosome reconstitution, 0.28 μM nucleosomes and 0.3 μM histones were incubated for 30 min at 20 °C in the low salt buffer. Samples were analyzed by native-PAGE in 0.5x TBE and visualized by ethidium bromide staining.

### Circular dichroism spectroscopy

Circular dichroism (CD) spectroscopy was performed on a J-815 CD Spectropolarimeter (Jasco). Spectra were measured in a range of 250-190 nm with 0.1 nm data intervals and 200 nm/min scanning speed. Proteins were analyzed in a quartz cuvette of 1 mm light path. Spectra were averaged from five scans. Histone H1 obtained from calf thymus (Calbiochem) served as a reference.

### Ubiquitylation assay

For ubiquitylation assays, 8.2 ng/μl E1 (UBA1), 8.3 ng/μl E2 (UBCH5B), and 25 ng/μl E3 (HUWE1, catalytic domain, aa 3228-4374/1147 aa) were incubated with 0.3 μg/μl Ub (Sigma-Aldrich) and 11 ng/μl H1.2/H1.2 KxAc in 2 mM ATP, 2 mM MgCl_2_, 1 mM DTT, 25 mM Tris-HCl (pH 7.4), 50 mM NaCl. Samples were incubated for 90 min at 30 °C and reactions stopped by the addition of SDS-PAGE loading buffer and heating to 95 °C for 5 min. Finally, samples were analyzed by SDS-PAGE and Coomassie blue staining.

### HEK 293T cell culture and lysate preparation

HEK 293T cells were cultured in DMEM supplemented with 10% (v/v) fetal bovine serum at 37 °C, 5% CO_2_. Cells were harvested by centrifugation, washed with ice-cold 1x PBS, and pellets were frozen in liquid nitrogen and stored at −80 °C.

Pellets were resuspended in 1x PBS, 2 mM MgCl_2_, 1 mM DTT, 100 μM Pefabloc SC, 1 μg/ml Leupeptin, 1 μg/ml Aprotinin, incubated on ice for 10 min and lysed by sonication. The lysate was cleared by centrifugation (30 min, 21,885 × g, 4 °C), and the protein concentration of the supernatant was determined by Bradford assay.

### H1.2 KxAc stability assay

To determine the stability of the acetylated H1.2-conjugates in human cell lysates, 4.7 μM H1.2 or H1.2 KxAc were incubated in HEK 293T cell lysates (5 mg/ml) at 4 °C or 25 °C for up to 24 h. Samples were analyzed by western blot with anti-H1 (Santa Cruz) and anti-AcK (Cell Signaling) antibodies. Experiments were performed in biological duplicates.

### Identification of the H1.2 KxAc interactome by AP-MS

To identify protein-protein interactions, pulldown assays were performed with H1.2 and H1.2 KxAc variants as described.^15^ Samples without bait protein served as control. 2.5 mg HEK 293T cell lysate and 2.35 nmol bait protein in 500 μl total volume were incubated on ice for 10 min and centrifuged for 10 min at 21,885 x g at 4 °C. The supernatant was added to 20 μl Strep-Tactin Superflow resin (IBA Lifescience) and incubated overnight at 4 °C. Beads were washed five times with 1x PBS, 2 mM MgCl_2_, 1 mM DTT, 100 μM Pefabloc SC, 1 μg/ml Leupeptin, 1 μg/ml Aprotinin followed by elution of bound proteins in 2.5 mM desthiobiotin. Samples were analyzed by SDS-PAGE (with 0.02% input and flow-through, 0.05% first washing fraction, 1.5% last washing fraction, 2.75% elution fraction loaded) and Krypton staining (Thermo Fisher Scientific). For identification of co-purified proteins, elution fractions were freeze-dried, followed by in-solution digestion. Samples were denatured in 8 M urea, reduced by the addition of 5 mM TCEP at 37 °C for 30 min, and subsequently alkylated in 10 mM iodoacetamide at room temperature for 30 min in the dark. After the addition of 50 mM NH_4_HCO_3_ to a final concentration of 1 M urea, samples were digested using trypsin (1:50, w/w, Promega) overnight at 37 °C. Finally, samples were acidified with TFA and lyophilized. Before LC-MS/MS analysis, samples were desalted using Micro-ZipTips (μ-C18, Merck).

Digested and desalted peptides were analyzed on an Orbitrap Fusion Tribrid mass spectrometer coupled to an EASY-nLC 1200 system (Thermo Fisher Scientific). Tryptic peptides were separated at a flow rate of 300 nl/min using a 190 min gradient from 5% ACN, 0.1% formic acid to 35% ACN, 0.1% formic acid and 10 min to 45% ACN, 0.1% formic acid followed by a washing step of 10 min at 80% ACN. MS spectra were recorded in the orbitrap at 120,000 (at m/z 200) resolution, scan range 300-1500 m/z, automatic gain control ion target value of 4e^5^, and maximum injection time of 50 ms. The intensity threshold was set to 5e^3^, exclusion duration to 45 s, and charge states to 2-7 were included. Fragmentation was performed by collision-induced dissociation (CID) with 35% collision energy and ions were subsequently detected in the ion trap. Automatic gain control was set to 2e^3^ with a maximum injection time of 300 ms. The system was operated in data-dependent top-speed mode with a 3 s cycle time. The affinity purification experiments were done in biological triplicates and each sample was measured twice as a technical duplicate.

Raw files from LC-MS/MS measurements were analyzed using MaxQuant 1.6.1.0^34, 35^ with match between runs and label-free quantification (minimum ratio count one) enabled. The minimal peptide length was set to five (further settings: default). For protein identification, the human reference proteome downloaded from the UniProt database (download date: 2018-02-22) and the integrated database of common contaminants were used. Further data processing was performed using Perseus 1.6.1.3 software^36^. Identified proteins were filtered for reverse hits, common contaminants, and proteins only identified by site. LFQ intensities were log2 transformed, filtered to be detected in at least four out of six replicates (i.e. three biological replicates each measured as technical duplicates), and missing values were imputed from a normal distribution (width = 0.3 and shift = 1.8 for total matrix). Significantly enriched proteins were identified by an ANOVA test (FDR = 0.01, s0 = 2), normalized by Z-scoring, and averaged. Proteins only enriched (Z-score > 0.3) in the control samples using beads only were removed and finally, the H1.2 and H1.2 KxAc-enriched proteins were analyzed by hierarchical clustering (Euclidean distance) and plotted as a heatmap. Identified proteins within specific clusters were additionally plotted in profile plots. Only proteins with a minimum Z-score of 0.3 were considered for further analysis (see also Supporting Data S1) and protein classification by function was performed using PANTHER 15.0^37^ focusing on the ‘Protein Class’.

### Western blot analysis

For orthogonal validation of identified interactions in HEK 293T cell lysate, samples resulting from the affinity purification assay were analyzed by western blot. Primary antibodies were directed against PAF1 (Bethyl Laboratories), ASCC2, ASH2L, CGBP/CXXC1, H1.2, SET, SIRT1, SUZ12 (Abcam), AEBP2 (Novus Biologicals), and CTR9 (antibodies-online).

### Deacetylation Assay

A plasmid containing the cDNA of SIRT1 (669 aa/aa 1-3 + 82-747) including a C-terminal His_6_-tag was transformed into *E. coli* BL21 (DE3). Cells were cultured at 37 °C in LB medium and 1 mM IPTG was added at OD_600_ = 0.6-0.8. Cells were harvested after 6 h. After cell lysis, SIRT1 was purified on a HisTrap FF column (GE Healthcare) followed by size-exclusion chromatography (HiLoad 16/600 Superdex 75 pg, GE Healthcare). Protein concentration was determined by BCA assay and SDS-PAGE followed by Coomassie blue staining.

For SIRT1-dependent deacetylation of H1.2 K17Ac and H1.2 K64Ac, 1.6 μM H1.2 KxAc and 10 nM-2 μM SIRT1 were incubated with 3 mM NAD^+^, 1 mM MgCl2 in 50 mM Tris-HCl (pH 8.0), 150 mM NaCl for 15 min at 30 °C. For time-dependent deacetylation of H1.2 K17Ac and H1.2 K64Ac, reactions containing 50 nM SIRT1 were incubated for the times indicated. Reactions were stopped by the addition of SDS-PAGE loading buffer and heating to 95 °C. Deacetylation was analyzed by western blot with anti-H1 (Santa Cruz), anti-AcK (Cell Signaling), and anti-His HRP conjugated (Sigma-Aldrich) antibodies. Experiments were performed in biological duplicates.

## RESULTS AND DISCUSSION

### Acetylation of linker histone H1

To expand the toolbox of site-specifically modified linker histones, we generated variants of H1.2 containing AcK at distinct positions (H1.2 KxAc) by genetic code expansion. We used an modified, orthogonal pyrrolysine tRNA synthetase (AcKRS)/tRNA_CUA_ pair originating from *Methanosarcina barkeri* to incorporate AcK in response to an amber stop codon (TAG/UAG).^29, 31^ To obtain insight into the effects of acetylation on H1.2 function, we chose two positions: K17 in the NTD (H1.2 K17Ac) and K64 in the GD (H1.2 K64Ac). These modification sites have been identified in earlier proteomic studies and particularly acetylation at K17 has been identified in most human somatic subtypes as well as in mice and chicken, indicating a high degree of conservation for this modification site.^38, 39^ In a previous study, we could also confirm the acetylation of endogenous H1.2 at position K17 and K64 in HEK 293T cells, the cell type used throughout this work (Figure 1).^15^ Moreover, to additionally evaluate the effect of di-acetylated H1, we also generated the doubly modified H1.2 with AcK incorporated at positions K17 and K64 (H1.2 K17K64Ac) (Figure 2A).

**Figure 1.**
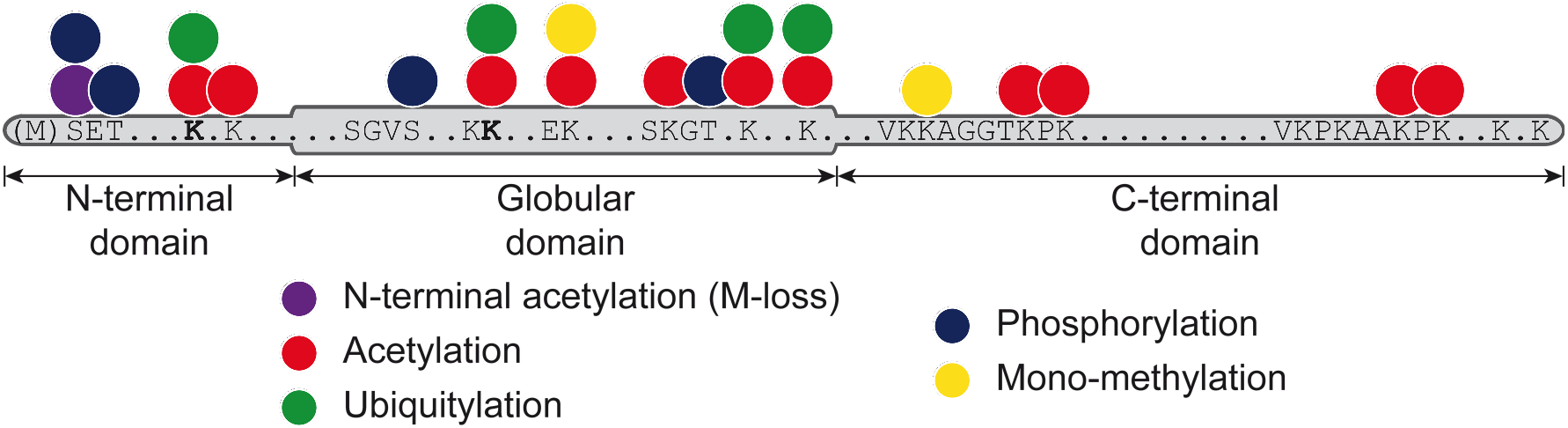
Modification sites of H1.2. Scheme of H1.2 PTMs after IP-enrichment from HEK 293T cells and identification by LC-MS/MS. Positions investigated within this study are indicated in bold (K17 and K64). Reprinted with adaptations from ^15^.

**Figure 2.**
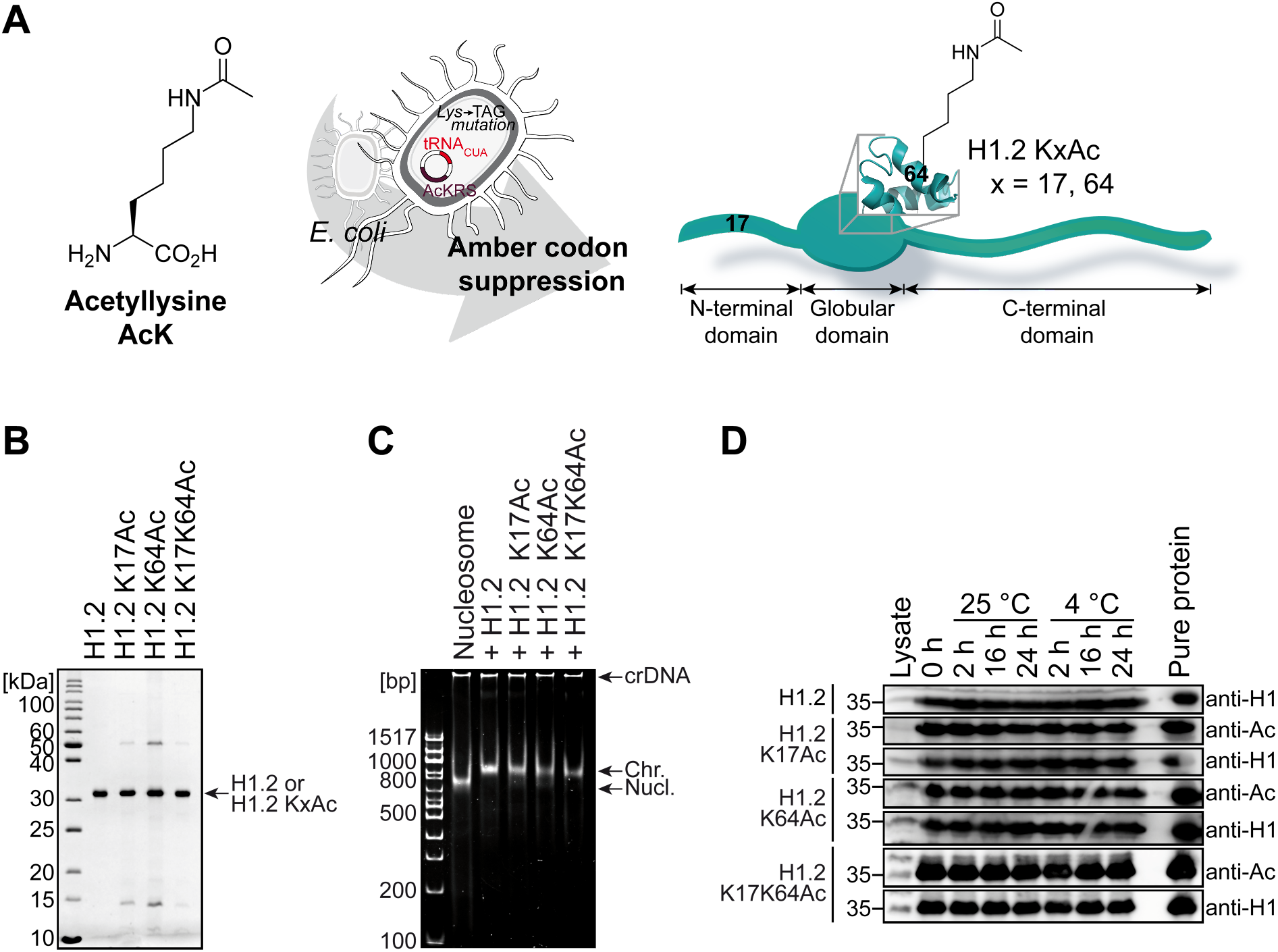
Generation of site-specifically acetylated H1.2. **(A)** AcK is incorporated into H1.2 during gene expression in *E. coli* in response to an amber stop codon (TAG/UAG) at positions K17, K64 (mono-acetylation), or both positions within one protein (di-acetylation). **(B)** Purified H1.2 and H1.2 KxAc variants resolved by SDS-PAGE and Coomassie staining. **(C)** Chromatosome assembly with H1.2 KxAc variants; crDNA indicates competitor DNA, Chr. refers to mono-chromatosomes, Nucl. indicates mono-nucleosomes. **(D)** Stability assay of acetylated histones in cell lysate. Proteins were incubated in HEK 293T cell lysate as indicated. Samples were analyzed by western blot showing constant levels of acetylated protein, indicating stability of the modification over time at different temperatures.

Site-specifically acetylated H1.2 variants were obtained in 0.5-0.9 mg yield per liter of bacterial culture (Figure 2B). LC-MS and LC-MS/MS analysis of the modified proteins demonstrated the homogeneous incorporation of one (H1.2 K17Ac and H1.2 K64Ac) or two (H1.2 K17K64Ac) AcK residues at the expected positions (Figure S1 and S2). The correct structural fold of the H1.2 KxAc variants was validated by CD spectroscopy (Figure S3). As shown in Figure 2C, binding of unmodified H1.2 and H1.2 KxAc variants to nucleosomes resulted in the formation of chromatosomes, further indicating their structural integrity. Moreover, using our histones as substrates in a ubiquitylation assay resulted in ubiquitylation of acetylated H1.2, demonstrating their acceptance as substrates by the ubiquitylation cascade (Figure S4). In summary, using genetic code expansion we could generate functionally viable, homogeneously and site-specifically mono- and di-acetylated H1.2.

### Identification of acetylation-specific H1 interactions

A prerequisite for the identification of PTM-specific interactions using cell lysates is that the modification is not removed by enzymes potentially present in the lysate. Thus, we next incubated the H1.2 KxAc variants with cell lysate and determined their acetylation status. Importantly, the acetylation status did not detectably change over time, indicating that no significant deacetylation takes place under the conditions used (Figure 2D). To determine whether site-specific acetylation affects the protein-protein interaction properties of H1.2, we incubated unmodified H1.2, mono-, and di-acetylated histones as well as empty beads with human HEK 293T cell lysate and identified interacting proteins by an AP-MS-based approach (Figure S5). In total, we identified 183 proteins that were consistently and significantly enriched over three biological replicate AP-MS experiments (ANOVA statistics, FDR = 0.01, s0 = 2; for a full list see Supporting Data S1).

To visualize the binding behavior of these proteins to the different baits, we applied hierarchical clustering resulting in the heatmap depicted in Figure 3. The heatmap can be subdivided into five clusters of proteins showing similar binding behavior as further indicated by their profile plots. Cluster 4 represents the largest fraction in which proteins bind to all H1.2 variants irrespective of their acetylation status. By contrast, clusters 1, 2, 3, and 5 show acetylation-dependent interactions with varying site-selectivity. Of these, cluster 1 includes mainly binders of H1.2 K17Ac and H1.2 K17K64Ac, and cluster 2 contains specific binders of all acetylated H1.2 variants. Cluster 3 shows interactors of H1.2, H1.2 K64Ac, and H1.2 K17K64Ac, whereas proteins of cluster 5 bind only unmodified and mono-acetylated H1.2.

**Figure 3.**
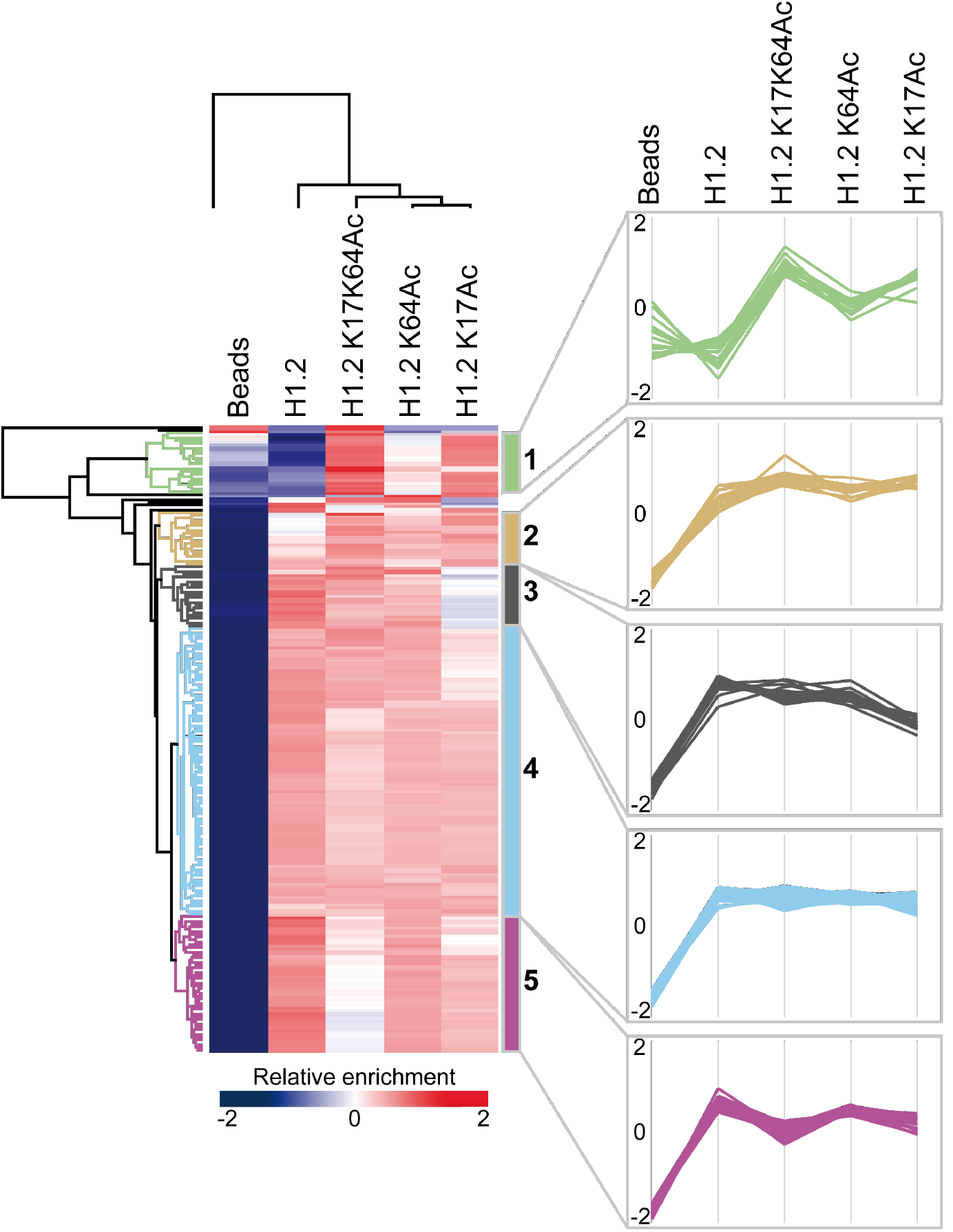
Identification of H1.2 KxAc-specific interactions. Heatmap representing the hierarchical clustering of statistically significant interactors following ANOVA analysis (FDR = 0.01, s0 = 2, n = 3). Interacting proteins are shown in rows, columns represent bait proteins. AP-MS experiments were carried out in biological triplicates with HEK 293T cell lysates. Empty beads (i.e. no bait protein) served as control. Clusters of proteins with similar interaction behavior are marked with different colors (left). Profile plots of clusters indicating specific binding patterns are shown on the right.

### Analysis of acetylation-specific H1 interactions

In the next step, we focused in more detail on the differential H1.2 KxAc-specific binders. Venn diagrams provide an overview of the acetylation site-specificity of the identified interactors (Figure 4A). While most protein interactors bind both unmodified and modified H1.2, 19% of the identified interactors were specifically enriched for acetylated histones only. More than 50% of these (i.e. 18 proteins) bind H1.2 K17Ac but not H1.2 K64Ac, indicating a distinct site-specificity for H1.2 K17Ac. Only three proteins exhibit site-specificity towards H1.2 acetylated at K64 (mono- and di-acetylation), while twelve proteins bind both H1.2 K17Ac and H1.2 K64Ac. Almost all interactors (32 of 33 proteins) of the mono-acetylated H1.2 variants bind di-acetylated H1.2, indicating that the two acetylation sites do not influence each other.

**Figure 4.**
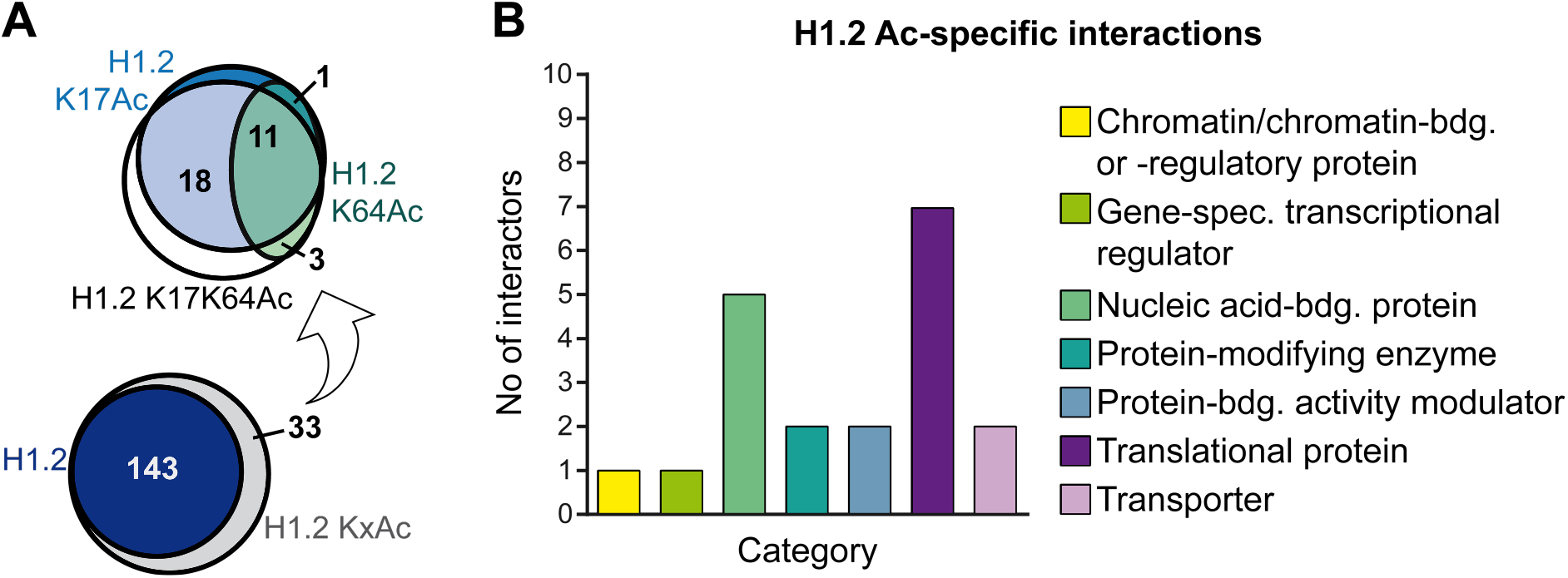
Acetylation-dependent interactions of H1.2 KxAc variants. **(A)** Venn diagrams of proteins specifically interacting with unmodified H1.2 and H1.2 KxAc (bottom), and acetylation-specific binders with site-specific resolution (top). **(B)** Gene Ontology analysis of all acetylation-specific H1.2 KxAc-interacting proteins based on PANTHER classification.

Gene Ontology (GO) analysis (Figure 4B) reveals that the acetylation-specific H1.2 interactors include, for example, translational proteins, such as five subunits of the EIF3 complex. Components of the EIF3 were already identified as binders of unmodified H1.0.^17^ Nucleic acid-binding proteins constitute another protein class enriched by H1.2 acetylation, including CTR9, a transcription factor and component of the PAF1 complex. Furthermore, we identified the deacetylase SIRT1, a chromatin and transcriptional regulator, as an H1.2 KxAc-specific interactor.

Interestingly, acetylation of H1.2 did not only result in the generation of new protein-protein interactions but also interfered with the binding of some interactors of unmodified H1.2 as 38% (only mono-acetylation) and 75% (mono- and di-acetylation) of unmodified H1.2 interactors showed reduced binding upon histone acetylation (Figure 3 and Supporting Data S1), indicating a major influence of the acetylated lysine residue on respective interaction sites.

In summary, the interactome data indicate that site-specific acetylation of H1.2 results in overlapping but distinct groups of interacting proteins.

To corroborate the results obtained by quantitative proteomics, the binding behavior of several proteins was analyzed by affinity purification followed by western blotting (AP-WB). Nine of the identified H1.2 interactors belonging to different clusters were selected (Figure 5). Comparison of enrichment patterns obtained from AP-WB and AP-MS results revealed highly similar interaction patterns, validating the approach in general and the accuracy of the acetylation-specific interactions as detected by interaction proteomics in particular.

**Figure 5.**
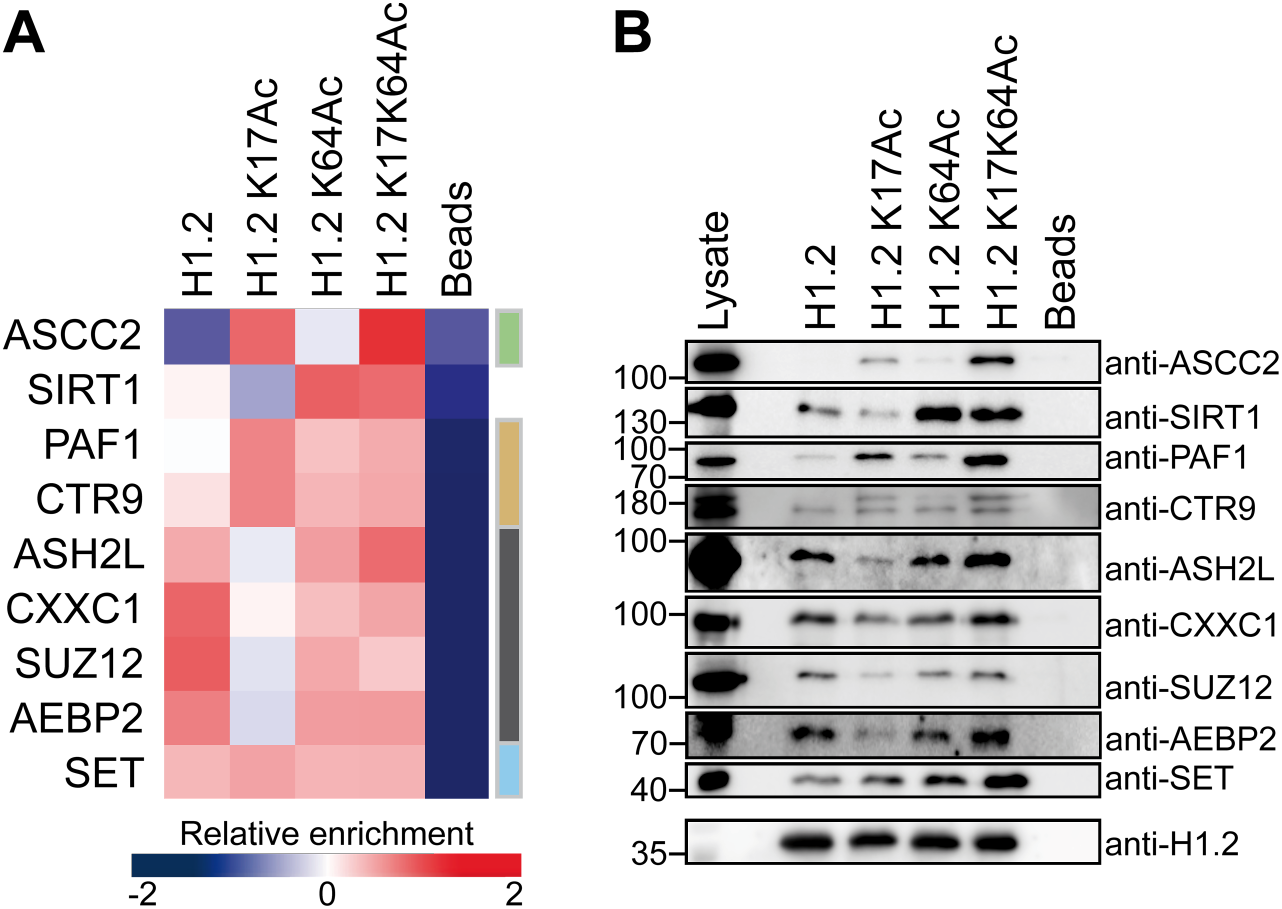
Validation of acetylation-specific interactions by western blot. **(A)** Heatmap showing relative enrichment of selected proteins as identified by AP-MS and **(B)** by immunoblotting. Lysate and elution fractions of the affinity purification assay were subjected to western blot analysis with antibodies specific for the respective proteins.

For example, the multitasking protein SET, which acts as a histone chaperone^40^ and is also involved in the inhibition of histone acetylation^41^, shows enrichment for all H1.2 variants studied. ASCC2, a subunit of the ASCC complex (activating signal cointegrator complex), which is involved in DNA damage repair,^42^ shows distinct specificity for H1.2 K17Ac and H1.2 K17K64Ac. This suggests a potential role of H1.2 acetylation at K17 in the DNA damage response.

PAF1 and CTR9 are subunits of the PAF1 complex, a transcription elongation factor and interactor of RNAPII (RNA Polymerase II). An and co-workers^16^ described the interaction of the PAF1 complex with H1.2 in combination with increased gene expression *via* ubiquitylation and methylation of the core histones. In our study, PAF1 and CTR9 show weak binding to unmodified H1.2, while both proteins are enriched for the H1.2 KxAc variants. This may indicate that acetylated H1.2 binds also stronger to the PAF1 complex than unmodified H1.2 and that H1.2 Ac is involved in transcriptional activation.

In contrast, H1.2 acetylated at position K17 exhibits reduced binding to SUZ12 and AEBP2, which are subunits of the histone methyltransferase PRC2 complex (polycomb repressive complex 2), as well as to ASH2L, a subunit of the SET1/MLL histone methyltransferase complexes, and CXXC1, a regulator of the SET1 complexes. While PRC2 catalyzes H3 K27 methylation leading to transcriptional repression,^43^ SET1/MLL histone methyltransferases are generally associated with transcriptional activation *via* methylation of histone H3 at position K4.^44, 45^ The reduced interaction of H1.2 K17Ac with subunits of these complexes suggests again a role of H1.2 Ac in modulating their activity, even though the opposite down-stream effects clearly warrant future investigations.

The NAD^+^-dependent protein deacetylase SIRT1 is involved in epigenetic regulation and has been identified as an interactor of H1.4 and H1.5. Furthermore, SIRT1 deacetylates H1.4 at position K26 and has been linked to the formation of facultative heterochromatin^21, 46^. We could recently show that site-specific ubiquitylation of H1.2 at position K64 also modulates interactions with SIRT1 and affects H1-dependent chromatosome assembly and phase separation.^15^ Here, we identified SIRT1 as mainly enriched with H1.2 K64Ac and di-acetylated H1.2 K17K64Ac, indicating a crucial role also for acetylation of H1.2 at position K64 for this interaction. *In vitro* assays show deacetylation activity of SIRT1 towards both positions studied, K17 and K64 (Figure 6). This confirms SIRT1 as a general deacetylase of H1 proteins and expands its scope to H1.2 K17Ac and H1.2 K64Ac *in vitro*. Taken together this indicates that both site-specific ubiquitylation and acetylation modulate the interaction of H1 with SIRT1 and suggests potential crosstalk of H1.2 acetylation and ubiquitylation.

**Figure 6.**
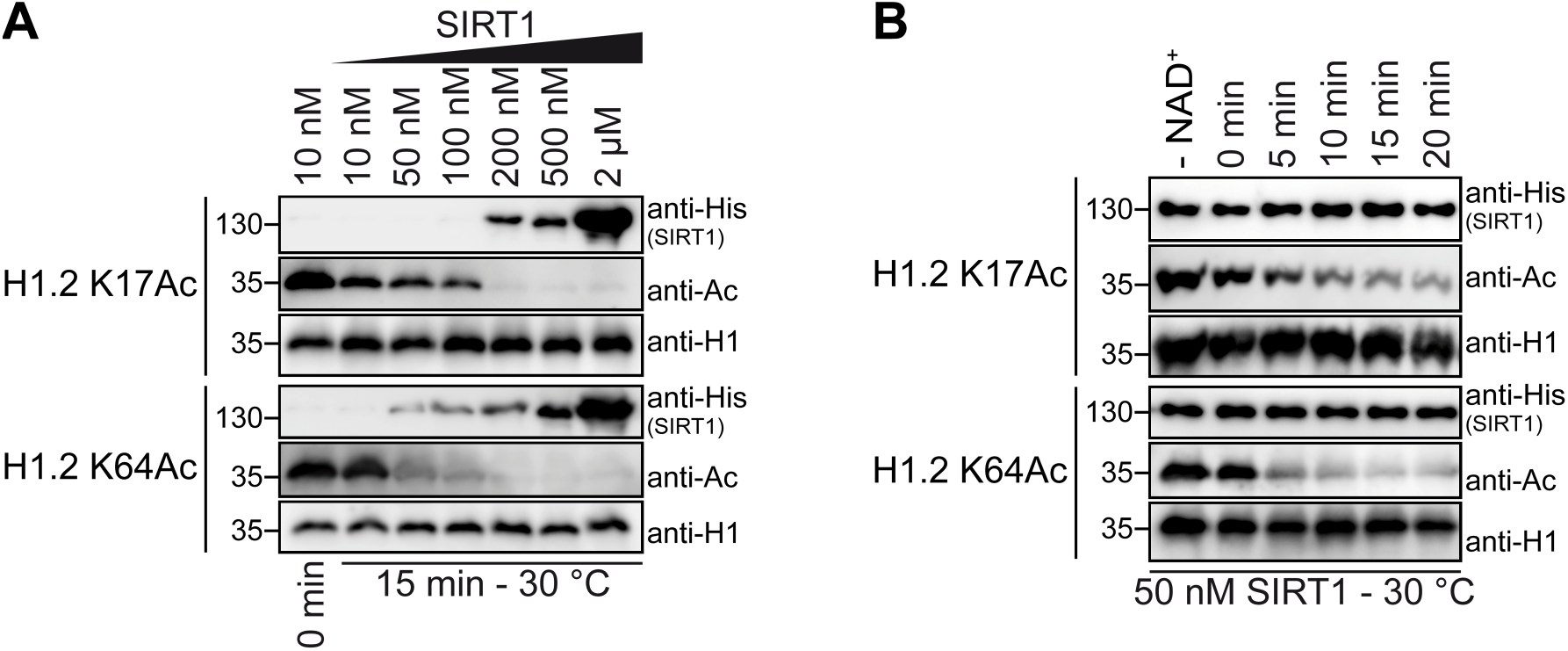
Deacetylation of the acetylated H1.2 using SIRT1. *In vitro* **(A)** concentration- and **(B)** time-dependent deacetylation of H1.2 K17Ac and H1.2 K64Ac by the NAD^+^-dependent deacetylase SIRT1. Protein and acetylation intensities were visualized by western blot with the indicated antibodies.

## CONCLUSIONS

Our results show the generation and application of site-specifically and homogeneously acetylated linker histones for the identification of acetylation-specific interactors and indicate that site-specific acetylation acts as a modulator of the H1 interactome.

## ASSOCIATED CONTENTS

### Supporting Information

Figures S1–5: LC-MS analysis of full-length proteins. LC-MS/MS analysis of acetylated histones. CD spectra of recombinant linker histones. Ubiquitylation assay using the H1.2 KxAc variants as substrate proteins. Identification of the H1.2 KxAc interactome by proteomic profiling.

### Supporting Data S1

Results of AP-MS analysis showing identified interactors.

### Data availability

The MS raw data have been deposited to the ProteomeXchange Consortium *via* the PRIDE^47^ partner repository with the dataset identifier PXD025832.

## Author Contributions

The manuscript was written through contributions of all authors. All authors have given approval to the final version of the manuscript. E.H., M.S., A.M., and F.S. conceived the study and experimental approach; E.H. and K.G. generated H1.2 and H1.2 KxAc variants with help from S.M.K. E.H. and K.G. performed AP-MS experiments. E.H., K.G., M.S., A.M., and F.S. analyzed the data, and E.H., M.S., A.M., and F.S. wrote the paper with input from all authors.

## Funding Sources

This work was supported by the DFG (SFB969). E.H. acknowledges the Konstanz Research School Chemical Biology for support by a fellowship. F.S. is grateful for funding from the DFG Emmy Noether Program (STE 2517/1-1).

## Notes

The authors declare no competing interests.

## ACKNOWLEDGMENTS

The authors thank the current and former members of the Marx, Stengel, and Scheffner labs for valuable discussions. We thank K. Stuber and F. Offensperger (Department of Biology, University of Konstanz) for providing purified E1/E2/E3 enzymes. We also thank A. Marquardt and A. Sladewska-Marquardt (Proteomics Center, University of Konstanz) for discussion.

## ABBREVIATIONS

Ac: acetylation
AcK: acetyllysine
AcKRS: pyrrolysine tRNA synthetase
ACN: acetonitrile
AP-MS: affinity purification-mass spectrometry
AP-WB: affinity purification followed by western blotting
CD: circular dichroism
CuAAC: Cu(I)-catalyzed azide-alkyne cycloaddition, click reaction
*E. coli*: *Escherichia coli*
H1.2 Ac: acetylated H1.2
H1.2 KxAc: H1.2 acetylated at lysine position x
LC-MS: liquid chromatography-mass spectrometry
LC-MS/MS: liquid chromatography-tandem mass spectrometry
PTM: posttranslational modification

## Supporting Information

**Figure S1.**
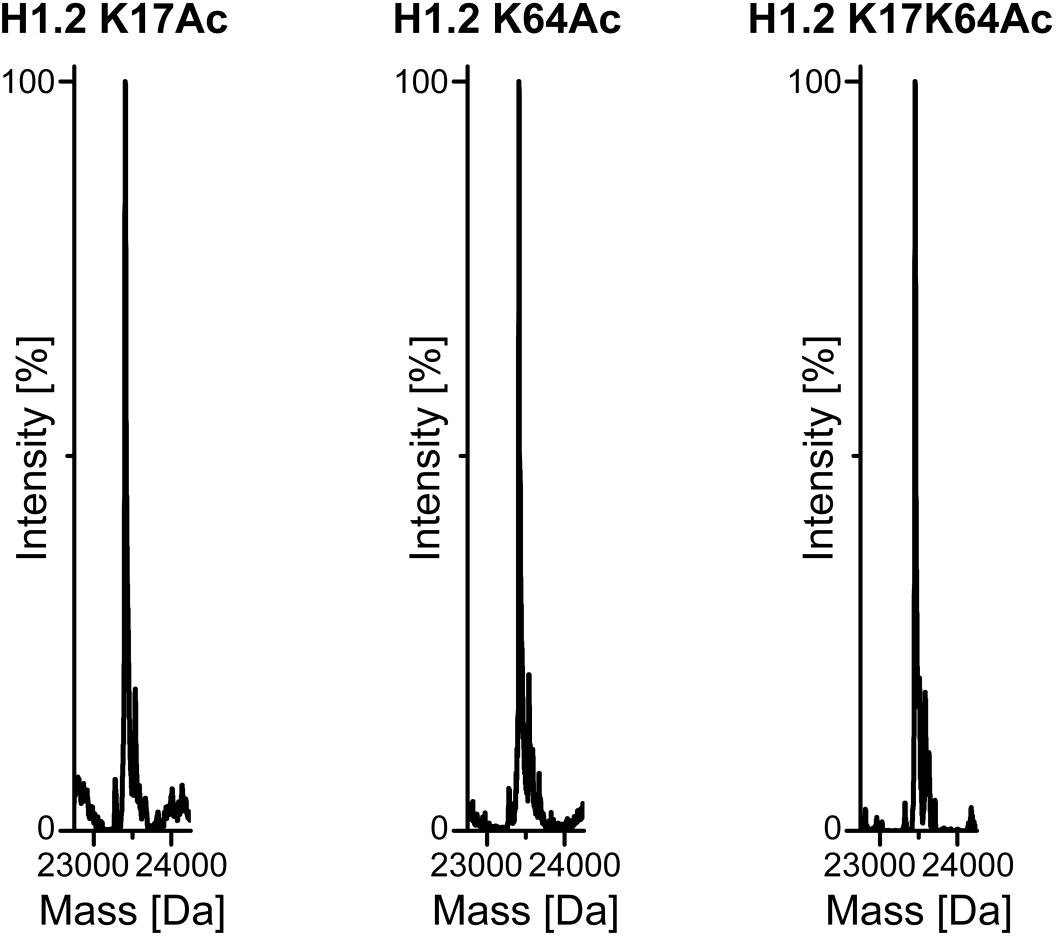
LC-MS analysis of full-length proteins. H1.2 K17Ac calc. 23412 Da, measured 23413 Da; H1.2 K64Ac calc. 23412 Da, measured 23413 Da; H1.2 K17K64Ac calc. 23454 Da, measured 23455 Da.

**Figure S2.**
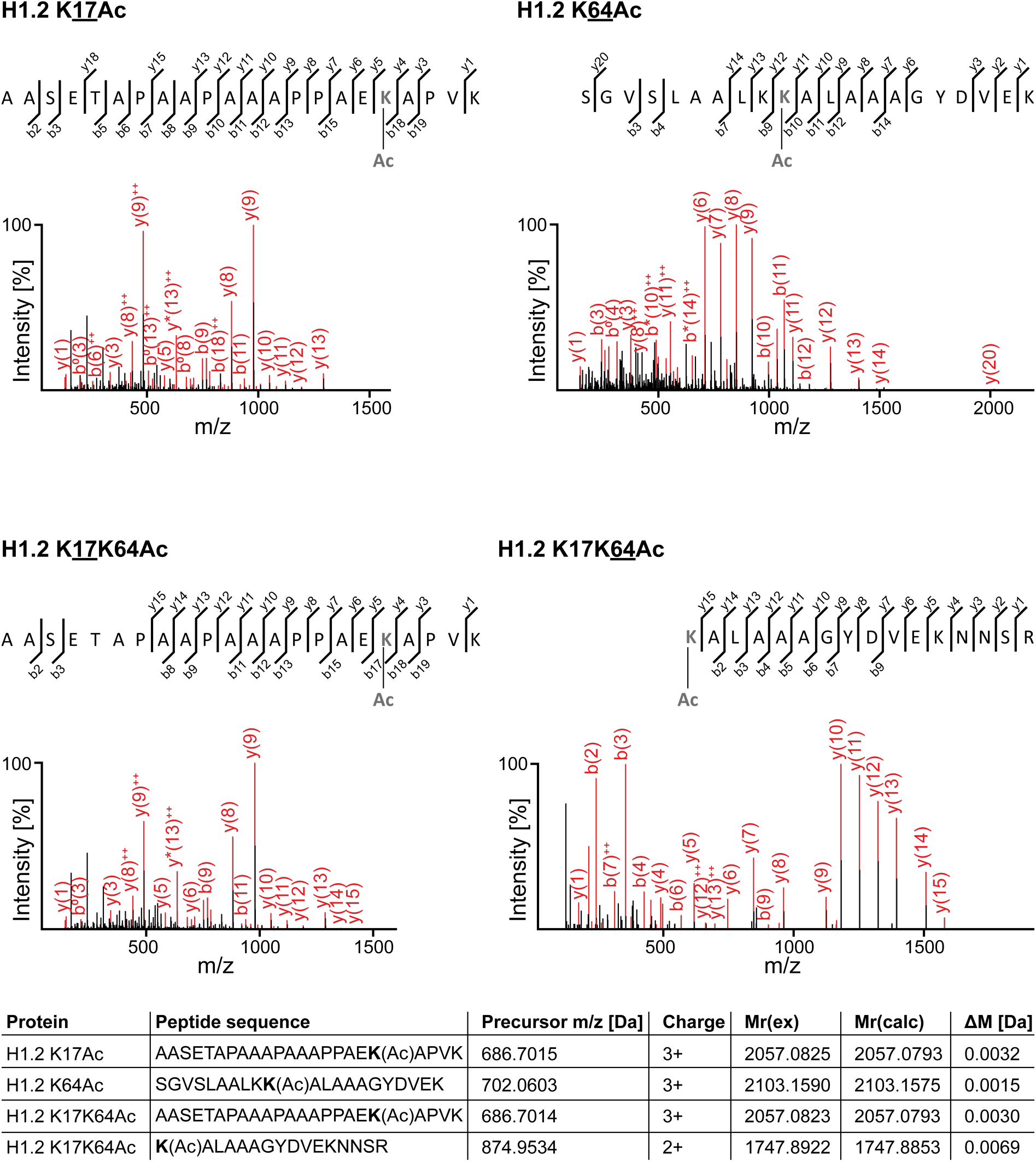
LC-MS/MS analysis of acetylated histones. Representative spectra showing incorporation of the non-canonical amino acid at the respective sites indicated with Ac. H1.2 K17Ac shown on the top left; H1.2 K64Ac on top right; H1.2 K17K64Ac shown in the middle, left position K17 and on the right position K64; the respective peptides are shown in the table depicted on the bottom.

**Figure S3.**
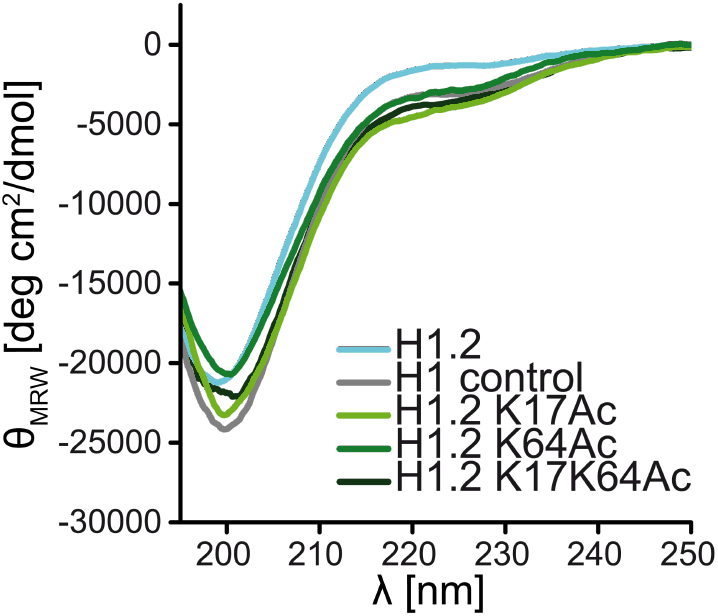
CD spectra of recombinant linker histones. CD spectra of unmodified H1.2, mono-acetylated H1.2, di-acetylated H1.2, and control H1 (Calbiochem) indicating correct protein refolding after purification under denaturing conditions independent of acetylation site.

**Figure S4.**
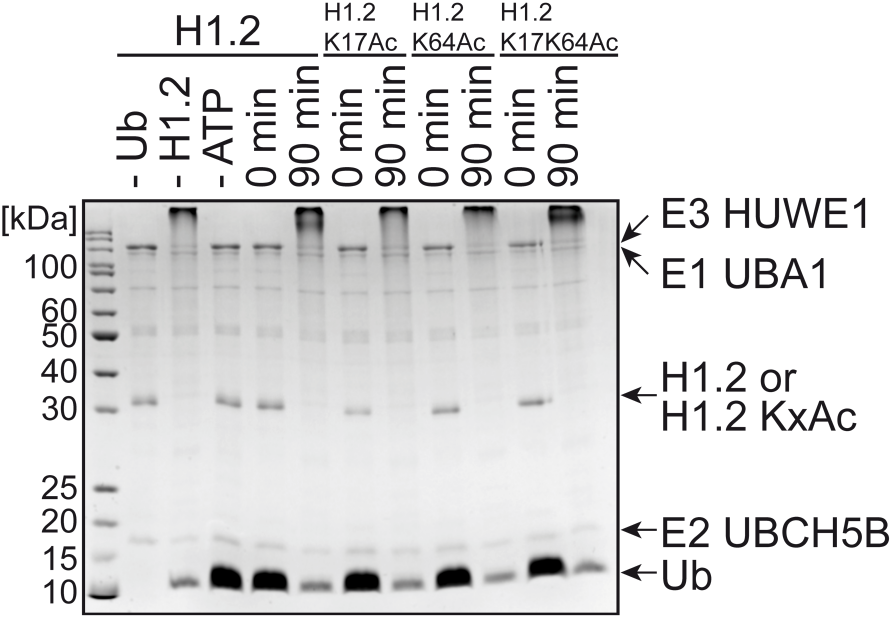
Ubiquitylation assay using the H1.2 KxAc variants as substrate proteins. *In vitro* ubiquitylation demonstrating the enzymatic acceptance of acetylated histones by the ubiquitylation cascade.

**Figure S5.**
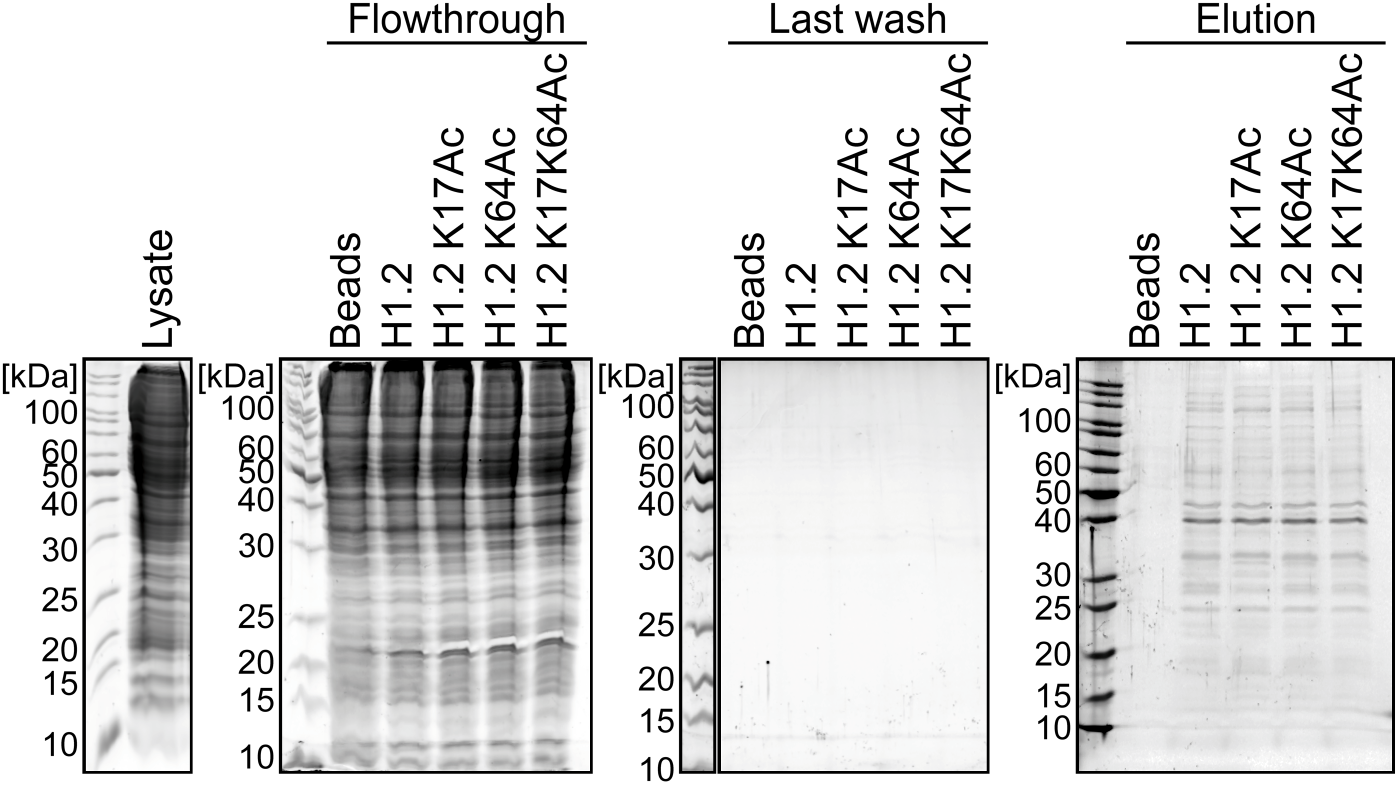
Identification of the H1.2 KxAc interactome by proteomic profiling. AP-MS assay analyzed by SDS-PAGE and Krypton protein staining. HEK 293T cell lysate was incubated with unmodified H1.2, H1.2 KxAc variants, and Strep-Tactin beads, or only empty beads as a control. Flowthrough was collected, the beads were washed five times, and bait proteins eluted with desthiobiotin along with the co-purified proteins.

## Notes

### Competing Interest Statement

The authors have declared no competing interest.

